# A representational asymmetry for composition in the human left-middle temporal gyrus

**DOI:** 10.1101/844183

**Authors:** Steven M. Frankland, Joshua D. Greene

**Affiliations:** Princeton Neuroscience Institute, Princeton, NJ; Harvard University, Cambridge, MA

**Author notes:** 33rd Conference on Neural Information Processing Systems (NeurIPS 2019), Vancouver, Canada.

## Abstract

Natural language is notable amongst representational systems for the rich internal structure of phrase and sentence-level expressions. Here, we provide evidence from two fMRI studies that a region of the left Middle Temporal Gyrus (MTG) exhibits a surprising representational asymmetry: verbs and patients (*to whom was it done?*) are bound to form a representation, but verbs and agents (*who did it?*) are not. Within MTG, BOLD signal to novel combinations of familiar components can be modeled by combining learned verb-patient conjunctive representations with more general agent representations, but not by the converse (verb-agent + patient). This asymmetry is not predicted by an abstract propositional representation of the event (e.g., chased(dog,cat), nor by a theory which derives conjunctions from the experienced statistical co-occurences between verbs and nouns. However, this asymmetry is predicted by various linguistic accounts of the internal structure of event descriptions (e.g., Williams, 1981; Marantz,1984; Grimshaw, 1990; Kratzer, 1996). These results provide evidence for the time-varying instantiation of re-usable representations of structure in MTG, consistent with the principle of compositionality, as well as accounts of verb-argument structure.

## 1 Background and Motivation

Theoretical models have long assumed that the data structures supporting the comprehension and production of human language are hierarchical and compositional. However, it does not follow that there must exist an isomorphism between the abstract, symbolic descriptions useful for linguistic analysis and the neural representations over which computations are performed. Here, we use functional magnetic resonance imaging (fMRI) to investigate the brain’s macro-level representational strategies for dynamically representing sentence structure and meaning.

A number of recent experiments have provided evidence that cortical regions surrounding the left sylvian fissure reflect aspects of compositionality (Pallier et al., 2011; Frankland and Greene 2015; Dehaene et al., 2015; Fedorenko et al. 2017; Nelson et al., 2017; Ding et al. 2017). Of course, such compositionality requires not only re-usable component representations, but the application of compositional operations to form representations with a particular *structure*, differentiating, for example, “the dog chased the cat” from “the cat chased the dog”. However, these representations may be considered *structured* at multiple levels that are not always distinguished. Here, we distinguish two levels of structure that are relevant to the construction of novel sentence meanings.

One level involves binding the same noun to different roles across instances (“the dog chased the man” vs. “the man chased the dog”). This level of structure, generally believed to involve “variable-binding”, has long been central to theoretical considerations in cognitive science (Fodor & Pylyshyn, 1988; Smolensky, 1990; Pinker 1995; Marcus, 2001; Kriete et al. 2013) as well as a motivation for recent empirical work in cognitive neuroscience (Frankland & Greene, 2015). However, linguistic analysis suggests the existence of additional non-obvious *asymmetric* structure in how these roles are treated in the grammar (Williams, 1981; Bresnan, 1982; Grimshaw, 1990; Marantz, 1984; Kratzer, 1996). It is generally agreed upon that some arguments are “internal” to the verb phrase (here, mapping on to the patient role in the event), while others are “external” to the verb phrase (here, the agent role), and, relatedly, that internal arguments are more closely tied to the interpretation of a verb’s meaning (Marantz, 1984). This difference between roles may owe to the order in which compositional operations are performed (Grimshaw, 1990), or to asymmetries in the lexical representation of the verb itself (Kratzer, 1996). Contrast this asymmetric representation with an abstract logical representation (e.g., chased(dog,cat)), which predicts *differences between nouns in particular roles* (dog-as-agent vs. dog-as-patient), but does not itself require any higher-order structural difference *between the roles themselves*, beyond difference in their identity.

In previous work (Frankland & Greene, 2015), we found that part of the left-mid superior temporal cortex (a) distinguishes between sentences comprised of the same parts (“the dog chased the man” vs. “the man chased the dog”) and (b) contains abstract agent and patient role representations that generalize across specific verbs. Here, we report analyses suggesting that a distinct part of the left-middle temporal gyrus, known to play a critical role in sentence processing (Dronkers et al. 2004, Hickok and Poeppel, 2007), exhibits key properties of compositionality that are consistent not only with the difference between nouns in a role, but also a higher-order asymmetry between roles.

## 2 Experiment 1

Here, we re-analyze data reported in experiment 2 of Frankland & Greene (2015). In this experiment, human participants (n=25) underwent high-resolution fMRI (1.5mm^3^ voxels, TR=2500 ms, TE=32 ms, 6/8th’s partial Fourier encoding, and 26 slices parallel to the AC-PC plane) while reading transitive sentences describing simple events comprised of an agent, event-type and patient (e.g., “the dog chased the cat”). The sentences were constructed using a menu of 4 nouns (“dog” / “cat” / “girl”/ “man”) and 5 verbs (“chased” / “scratched”/ “blocked”/ “approached”/ “bumped”). These items were re-combined to generate every possible agent-verb-patient combination, with the exception of self-combinations (e.g., participants never saw “the dog chased the dog”). Each of the 60 unique propositions (4X5X3) was presented once per scan run for 6 runs. Whether a particular sentence token was presented in the active or passive voice was randomly determined. For present purposes, active and passive versions of the same semantic relation were treated as equivalent (e.g., “chased by the dog” and “the dog chased” were coded identically).

In the present work, we focused on a left temporal ROI that contained parts of inferior, middle, and superior temporal gyrus (See Frankland & Greene, 2015). Within this ROI, we performed a searchlight analysis in which separate linear discriminant classifiers with a shrinkage estimate of the covariance matrix (Pereira &Botvinick, 2011) were trained to identify *who did it?* and *to whom was it done?* for a particular sentence, with scan runs treated as cross-validation folds. Critically, for the present analysis, the training and testing sets were iteratively restricted to include only one of the 5 verbs (e.g., chased), enabling the classifier to learn to map from patterns of activity to particular verb-role conjunctions. That is, one classifier would learn functions to perform the 4-way discrimination of *dog-as-chasee* from *man-as-chasee* and *cat-as-chasee* and *girl-as-chasee*, while a separate classifier would attempt to discriminate *dog-as-chaser*, from *man-as-chaser* etc. The classifier weights were then setparately re-learned and tested for each verb. The results were averaged across the 5 verbs (and 6 cross-validation iterations per verb) to generate one accuracy map for verb-patient classification and verb-agent classification for each subject. These maps were then warped to Talairach space and submitted to a second-level t-test against chance classification performance (25%). We tresholded searchlight results at p<0.01 voxelwise and corrected for multiple comparisons within the left temporal ROI, using clusterwise correction in AFNI.

This searchlight reveals a cluster within the left-MTG that carries information about verb-patient combinations (k=262, p<0.001 clusterwise corrected in left temporal ROI), but no significant clusters within the temporal lobes for verb-agent conjunctions. See Figure 1B. We then iteratively localized this verb-patient ROI in MTG by leaving each subject out of the localization procedure, and averaging the two search maps for the held out subject. Within these independently localized ROIs, within-verb patient classification was significantly better than within-verb agent classification (t(24)=2.5, p=0.02) and within-verb agent classification was not significantly better than chance (t(24)=0.59, p=0.28). Notably, there is no reason to expect such a difference for regions encoding the basic semantic content of the variables (e.g., dog vs. man). Anatomically, this region is centered in the lateral middle temporal gyrus (center, -61, -19, -4, Talairach). This differs from a nearby region of STS/STG in which the learned patterns generalize *across verbs* (See Figure 1c). In previous work, (Frankland & Greene 2015), we trained the classifier on a subset of verbs, and evaluated its ability to generalize the learned representations (e.g., man-as-agent) to novel verbs (e.g., “chased”). The *across-verb* (STG) and *within-verb* (MTG) patient ROIs differed significantly in their informational content (F(1, 24)= 8.35, p=0.008) when independently localized.

**Figure 1:**
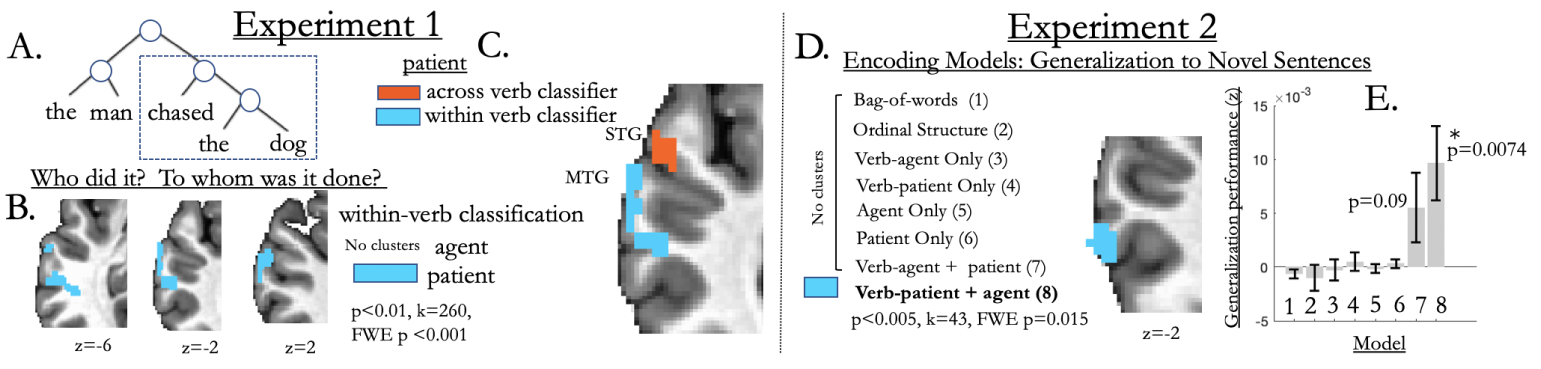
Two experiments suggest an asymmetry in MTG between the representation of agents and patients, consistent with a hierarchical syntactic structure, like that shown in (A). In experiment 1, (B) we find that a region of left-middle temporal gyrus (MTG) carries information about the patient (to whom was it done?) of an event, but not the agent (who did it?), when the classifiers are trained to learn verb-specific role-filler bindings (who was the chaser? who was the chasee?). (C) This representational profile differs significantly from a nearby region of superior temporal gyrus (STG) that carries information about the patient that is invariant across a class of verbs (e.g., representing ‘dog’ as the patient of ‘chased’/’scratched’/’bumped’ etc.). (D). In a second experiment, we find that a nearby region of MTG can be predicted as a linear combination of a learned verb-patient representation plus a general agent representation, but not the converse (verb-agent + patient).This is consistent with a particular abstract order of composition aligned with the deep syntactic structure shown in (A), in which the verb and patient are conjoined, and this conjunctive representation is then integrated with the agent.

## 3 Experiment 2

In experiment 1, we find that a region of left MTG carries verb-specific information about the patient, but not about the agent, even though the fillers were selected from the same menu (e.g., “the dog chased the cat”/ “the cat chased the dog”). In experiment 2, we attempt to predict BOLD signal in MTG as participants read *novel propositions* that are composed of familiar components. Here, a distinct group of participants (n=48) underwent fMRI (2mm^3^ voxels, TR=3500 ms, TE=28 ms, parallel imaging iPAT=2) while reading transitive sentences describing simple events comprised of an agent, event-type and patient. Here, sentences were generated from a menu of 6 nouns and 8 verbs creating 240 (6×8×5) unique propositions. Unlike experiment 1, each proposition was only presented once and was likely to be unfamiliar to participants (e.g., “the moose chased the cow”), as no active or passive sentence tokens were returned via Google search at the time the experiment was conducted.

We compared simple encoding models that predict BOLD signal at each voxel as a weighted linear combination of various sentence components. (See Frankland & Greene, (2019) for a more complete procedural description). Here, we compare encoding models based on bag-of-words, superficial ordinal structure (i.e., surface syntax), as well as underlying semantic variables shared across voices. This included variables encoding the agent or patient, dummy coded to involve one term per noun-role combination (e.g., moose-as-agent, cow-as-agent, moose-as-chaser, cow-as-chaser etc.) Critically, we also included models that predicted BOLD signal as a *combination* of multiple learned representations. One such model composed the verb-patient and general agent parameters. Another model composed learned verb-agent parameters with a general patient representation. The former is aligned with the hierarchical syntactic structure of the deep syntax, shown in Figure 1A, but the latter is not. Given that the number of training observations (200) always exceeded the number of model parameters (maximum=54) the beta values could be obtained analytically as least square estimates. To evaluate each model, we generate predictions using that model for each of the 40 held-out observations for each cross-validation iteration, and created a 40×40 matrix of the squared difference between every mapping of model-prediction and observation. The diagonal elements of this matrix represent the correct mapping between predictions and observations, and off-diagonal elements represent the incorrect mappings. We ask whether the squared difference is significantly better for the on-diagonal (match) than off-diagonal elements (mis-match) elements.

Notably, we find that the model using verb-patient + agent representations yields a significant cluster (p<0.005 voxelwise, k=43, p=0.015 FWE, small-volume corrected in *a priori* left temporal ROI), partly overlapping, but centered posteriorly to, the region of MTG found in experiment 1 (See Figure 1D). None of the remaining models significantly predict BOLD signal to novel sentences in MTG, including a verb-agent + patient representation. When this ROI is localized in independent subjects, the combined verb-patient+agent model is better than either the verb-patient (t(48)=2.9,p=0.006) or agent (t(48)=2.83, p=0.007) models in isolation (See Figure 1E). However, a direct contrast between the verb-patient + agent (syntax aligned) and verb-agent + patient (syntax-misaligned) models is not significant (t(48)=1.26, p=0.21). Given the results of experiment 1, it is somewhat surprising that we do not also see a significant cluster for the verb-patient model alone in MTG. One possibility is that the classification performance in Experiment 1 relies on pooling signals distributed across the region, which are ignored in the voxelwise encoding model. Better understanding the differences between these two analysis methods is an important topic for future work. Still, it is notable that the best model contains terms encoding the verb-patient interaction combined with terms for a more general agent representation. This is *prima facie* consistent with the hierarchical syntactic structure presented in Figure 1a.

## 4 Discussion

Taken together, these two studies suggest a higher-order asymmetry between the representation of the agent and patient roles in the left middle temporal gyrus (MTG). In previous work (Frankland & Greene, 2015), we found evidence that parts of MTG, STS, and STG carry information about *who did what to whom* in a sentence (Frankland & Greene, 2015), differentiating instances in which the same nouns are bound to different roles (e.g., ‘the dog chased the man’ vs. ‘the man chased the dog’). Although the physical mechanisms for variable-binding remain unclear, this high-level signature reflects this binding, however it occurs. Moreover, within an anterior and superior portion of lmSTC in the superior temporal gyrus, these role representations are shared across verbs. Here, we add to this work, finding that a nearby left temporal region (in MTG) is not only sensitive to the content of dynamic noun-role bindings, but also exhibits a higher-order asymmetry between thematic roles, encoding verb-patient conjunctions, but not verb-agent conjunctions.

Although neither a propositional representation (chased(dog, cat)) nor the statistics of the experiment (dog –> chased –> cat) would predict the observed asymmetry, it is expected if this particular region is tracking either the argument structure of the verb, or if it is mapping onto a latent hierarchical syntactic representation (See Marantz,1984; Grimshaw, 1990; Kratzer, 1996 for possibilities). We are agnostic on the proper theoretical construal of this asymmetry. One possibility is that this reflects the abstract (though not necessarily sequential) order of compositional operations, whereby the object (patient) and verb are integrated in the verb-phrase, and then combined with the subject (agent). Another possibility is that these MTG codes reflect the saturation of the arguments retrieved from the lexical representation of the verb, which may only have an internal argument, and no external argument (Kratzer, 1996). Whatever the proper construal, our evidence suggests that two regions of the left temporal lobes correspond to different levels of representation relevant to mapping between event representation and sentence syntax: STG/STS represents *abstract* role representations, recurring across verbs, as in the classic semantic roles (Gruber, 1965; Parsons, 1990). MTG contains verb-specific roles that asymmetrically link the verb and patient, but not the verb and agent (Marantz, 1984; Kratzer, 1996). At a high-level, we expect the syntactic functions that take internal or external arguments to be parameterized by semantic relations in the event representation (e.g., Williams, 1981). This may be supported by bi-directional interactions between the distinct representations of structure in MTG and STG.

## References

Bresnan, J. (1982). The mental representation of grammatical relations (Vol. 1). The MIT Press.

Dehaene, S. Meyniel, F. Wacongne, C. Wang, L. & Pallier, C. (2015). The neural representation of sequences: from transition probabilities to algebraic patterns and linguistic trees. Neuron, 88(1), 2–19.

Ding, N. Melloni, L. Zhang, H. Tian, X. & Poeppel, D. (2016). Cortical tracking of hierarchical linguistic structures in connected speech. Nature neuroscience, 19(1), 158.

Dronkers, N. F., Wilkins, D. P., Van Valin Jr, R. D., Redfern, B. B., & Jaeger, J. J. (2004). Lesion analysis of the brain areas involved in language comprehension. Cognition, 92(1-2), 145–177.

Fedorenko, E. Scott, T. L. Brunner, P. Coon, W. G. Pritchett, B. Schalk, G. & Kanwisher, N. (2016). Neural correlate of the construction of sentence meaning. Proceedings of the National Academy of Sciences, 113(41), E6256–E6262.

Fodor, J. A., & Pylyshyn, Z. W. (1988). Connectionism and cognitive architecture: A critical analysis. Cognition, 28(1-2), 3–71.

Frankland, S.M. & Greene, J.D. (2015). An architecture for encoding sentence meanings in left-mid superior temporal cortex. Proceedings of the National Academy of the Sciences, 112(37), 11732–11737.

Frankland, S.M. & Greene, J.D. (2019). Distinct forms of compositional semantic representation across brain regions. PsyArxiv.

Grimshaw, J. (1990). Argument structure. MIT Press. Cambridge, MA.

Gruber, J. S. (1965). Studies in lexical relations (Doctoral dissertation, Massachusetts Institute of Technology).

Hickok, G., & Poeppel, D. (2007). The cortical organization of speech processing. Nature reviews neuroscience, 8(5), 393.

Kratzer, A. (1996). Severing the external argument from its verb. In Phrase structure and the lexicon. pp. 109–137. Springer, Dordrecht.

Kriete, T., Noelle, D. C., Cohen, J. D., & O’Reilly, R. C. (2013). Indirection and symbol-like processing in the prefrontal cortex and basal ganglia. Proceedings of the National Academy of Sciences, 110(41), 16390–16395.

Marantz, A. (1984). On the nature of grammatical relations. MIT Press: Cambridge, MA.

Marcus, G. F. (2001). The algebraic mind: Integrating connectionism and cognitive science. MIT press: Cambridge, MA.

Nelson, M. J. El Karoui, I. Giber, K. Yang, X. Cohen, L. Koopman, H. & Dehaene, S. (2017). Neurophysiological dynamics of phrase-structure building during sentence processing. Proceedings of the National Academy of Sciences, 114(18), E3669–E3678

Pallier, C. Devauchelle, A. D. & Dehaene, S. (2011). Cortical representation of the constituent structure of sentences. Proceedings of the National Academy of Sciences, 108(6), 2522–2527.

Parsons, T. (1990). Events in the Semantics of English (Vol. 334). Cambridge, MA: MIT press.

Pereira, F., & Botvinick, M. (2011). Information mapping with pattern classifiers: a comparative study. Neuroimage, 56(2), 476–496.

Pinker, S. (1997). How the mind works. Penguin: UK.

Smolensky, P. (1990). Tensor product variable binding and the representation of symbolic structures in connectionist systems. Artificial intelligence, 46(1-2), 159–216.

Williams, E. (1981). Argument structure and morphology. The linguistic review, 1(1), 81–114.

